# Multi-objective optimisation of material properties and strut geometry for poly(L-lactic acid) coronary stents using response surface methodology

**DOI:** 10.1101/667915

**Authors:** R.W. Blair, N.J. Dunne, A.B. Lennon, G.H. Menary

## Abstract

Coronary stents for treating atherosclerosis are traditionally manufactured from metallic alloys. However, metal stents permanently reside in the body and may trigger undesirable immunological responses. Bioresorbable polymer stents can provide a temporary scaffold that resorbs once the artery heals but are mechanically inferior, requiring thicker struts for equivalent radial support, which may increase thrombosis risk. This study addresses the challenge of designing mechanically effective but sufficiently thin poly(L-lactic acid) stents through a computational approach that optimises material properties and stent geometry. Forty parametric stent designs were generated: cross-sectional area (post-dilation), foreshortening, stent-to-artery ratio and radial collapse pressure were evaluated computationally using finite element analysis. Response surface methodology was used to identify performance trade-offs by formulating relationships between design parameters and response variables. Multi-objective optimisation was used to identify suitable stent designs from approximated Pareto fronts and an optimal design is proposed that offers comparable performance to designs in clinical practice. In summary, a computational framework has been developed that has potential application in the design of high stiffness, thin strut polymeric stents that contend with the performance of their metallic counterparts.

## 1. Introduction

Balloon angioplasty, performed by Andreas Grüntzig in 1977, is recorded as the first successful effort to treat an occluded coronary artery and subsequently revolutionised the treatment of coronary artery disease.^[1]^ However, the surgical procedure suffers from significant limitations, namely vessel occlusion and restenosis, which prompted the development of the first bare metal stent *(BMS)* nearly a decade later.^[2]^ Whilst *BMSs* reduced the incidence rate of restenosis when compared to balloon angioplasty, the introduction of a permanent metallic cage provoked neointimal hyperplasia, an inflammatory response of the vessel walls^[3]^, and as a result drug-eluting stents (*DESs*) succeeded *BMSs*, containing a durable polymer coating which releases an antiproliferative drug (e.g. sirolimus or paclitaxel) that attenuates intra-stent neointimal proliferation^[4]^. Drug-eluting stents have shown reduced restenosis rates when compared to *BMSs*.^[5,6]^ However, they suffer from inherent flaws based on the permanent nature of their design and issues have been reported regarding the long-term (> 1 year) safety of these devices including delayed healing and late stent thrombosis (*LST*)^[7,8,9]^, which has prompted the development of bioresorbable stents (*BRSs*). Bioresorbable stents provide short-term scaffolding to the arterial wall until it has healed and are subsequently resorbed, offering superior conformability and flexibility to their permanent metallic counterparts, whilst enabling late luminal gain, late expansive remodelling and potentially reducing the risk of *LST* associated with *DESs* following resorption.^[10,11]^

Whilst polymeric *BRSs* present a clinically attractive option, they require wider and thicker struts to provide an equivalent level of arterial support (Table 1) when compared to their metallic counterparts. As a result, polymeric *BRSs* have higher stent-to-artery ratios,^[12,13]^ which have been shown to increase the risk of myocardial infarction, thrombosis and restenosis.^[14,15]^ A thick-strut design also limits the diameter a stent can be crimped to, resulting in an increased crossing-profile that hinders the deliverability of the device^[16]^ and restrict normal vasomotion.^[4]^ Additionally, polymeric *BRS*s demonstrate higher degrees of foreshortening (due to an increased strut length) during deployment, which can initiate vascular restenosis injuries.^[17]^ Improvements in material processing, coupled with the correct matching of the stent geometry to the material may produce polymeric *BRSs* with reduced strut thickness and comparable performance to current generation metallic *DES*.^[18–20]^

**Table 1.** Comparison of strut geometry and performance metrics of clinically tested bioresorbable stents (*BRSs*) and modern metallic drug-eluting stents (*DESs*) for coronary application.^[4,12,20–25]^

|  | *Polymeric BRSs* | *Metallic DESs* |
| --- | --- | --- |
| Strut thickness ($\mu\text{m}$) | 125–156 | 80–140 |
| Strut width ($\mu\text{m}$) | 140–216 | 80–132 |
| Stent-to-artery ratio (%) | 26.0–32.0 | 15.5–21.4 |
| Crossing profile (mm) | 1.2–1.7 | 1.0–1.2 |

The elastic modulus of the polymer, which affects the radial collapse pressure of the stent, may potentially be the most important parameter in polymeric *BRS* design.^[20,26]^ Pauck and Reddy^[20]^ performed computational bench testing on three commercially available stent geometries, whilst varying the elastic modulus of the platform material, poly(L-lactic acid) (*PLLA*). The authors concluded that using a geometry similar to that of the Absorb *BVS* (Abbott Vascular, USA), with a strut thickness and a strut width of 100 μm, coupled with an elastic modulus of 9 GPa, allows the desired collapse pressure of at least 40 kPa to be met.^[18]^ The elastic modulus of extruded *PLLA* is approximately 3 GPa,^[27]^ which is significantly lower than the required value of 9 GPa, and hence additional processing steps must be taken to improve upon this.

Stretch blow moulding (*SBM*) is a processing technique used in the production of *BRS* to improve the elastic modulus of the polymer.^[27,28]^ In the SBM process, the polymer is initially extruded into a thick-walled tube (parison) and heated above its glass transition temperature during which it is biaxially stretched to create a thin-walled tube with improved mechanical properties.^[29]^ Whilst a three-fold increase in the elastic modulus is difficult to physically attain, Blair et al.^[30]^ showed that by tailoring processing parameters, biaxial stretching can improve the elastic modulus and yield strength of extruded *PLLA* sheet by approximately 80% and 70%, respectively. Given that the relationship between elastic modulus and strut thickness has been shown to be nonlinear,^[26]^ through careful matching of material properties to stent geometry, a physically attainable elastic modulus may be used to meet the radial stiffness threshold with a minimal increase in strut thickness.

The mechanical performance and efficacy of a stent design is strongly dependent on the configuration of strut geometry.^[31–33]^ Finite element analysis is an especially prevalent technique within the discipline of computational biomechanics, where *in vivo* testing is exceptionally challenging, and may be used as preclinical testing tool to optimise stent geometry prior to any form of physical testing.^[31,34]^ To evaluate the performance and efficacy of a given stent design, simulated tests are typically conducted in which one (or more) metrics are assessed across a range of potentially viable stent geometries. Stent geometries may be parameterised in terms of strut width, strut thickness, strut length and connector shape^[35]^ whilst performance metrics fall under two main headings: (i) dilation metrics and (ii) mechanical metrics. Dilation metrics are concerned with the behaviour of the stent during (and immediately following) inflation, with radial recoil, foreshortening and stent-to-artery ratio amongst the most commonly evaluated metrics.^[32,36]^ Mechanical metrics are concerned with the performance of the expanded stent, with radial stiffness considered as the most important mechanical metric for polymeric stents.^[20]^

It is difficult to define what constitutes an optimal stent design, given that the definition of ‘optimal’ depends on the parameters investigated and the performance metrics assessed. The ideal stent is typically considered as one that is highly deliverable with thin-struts (to improve delivery through tortuous vascular paths) but with high radial stiffness and minimal elastic recoil, to resist restenosis.^[37]^ However, this statement in itself presents a number of conflicting requirements and as a result, an optimised design will always be a trade-off. This is evident from a cross-comparison of the parametric studies conducted by García et al.,^[38]^ Li et al.,^[39]^ Migliavacca et al.,^[32]^ Pant at al.^[40]^ and Timmins et al.^[41]^ Radial stiffness and radial recoil were improved by increasing strut width and strut thickness whilst decreasing strut length, however this often came at the expense of the stent-to-artery ratio and foreshortening.

In summary, improvements in *PLLA* stent design may be attained using a combination of two factors: (i) enhancing mechanical properties of the platform polymer by tailoring its processing history and (ii) iteratively refining the stent’s shape by modifying key geometric features. Few studies have considered the combined effect of the processing history and stent geometry in order to optimise stent performance.^[39,42]^ Furthermore, to the best of the authors’ knowledge, no study has considered the combined effect of the biaxial stretching processing history and the geometric configuration when optimising the mechanical performance of a coronary stent. This study aims to address this challenge of designing mechanically effective but sufficiently thin bioresorbable *PLLA* stents through multi-objective optimisation of material parameters and stent geometry.

## 2. Material and methods

The design of *PLLA* stents may be improved by enhancing the material properties of the platform polymer through biaxial stretching and iteratively refining the stent geometry. By parameterising these design inputs and computationally evaluating the performance of a given stent design across a series of metrics (that capture the conflicting requirements for a stent), empirical relations were established that relate both the stent processing history and geometry to its performance. Using these empirical relations, performance trade-offs were identified and an optimal design may be identified through multi-objective optimisation.

### 2.1 Process parametrisation

In a previous study by Blair et al.^[30]^, the *SBM* process was idealised and replicated using a custom-built biaxial tensile tester, to evaluate the mechanical properties of *PLLA* pre- and post-biaxial stretching. The elastic modulus (*E*) and yield strength (*σ_Y_*) of extruded PLLA sheet increased by approximately 80% and 70% following biaxial stretching. These mechanical properties were observed to be highly dependent on the stretch ratio in the machine direction (*MD*), *λ_MD_*, and the stretch ratio in the transverse direction (*TD*), *λ_TD_*, in addition to the aspect ratio (*A_r_*) between the pair, defined as the quotient of *λ_TD_* and *λ_MD_* (Fig. 1).

**Fig. 1.**
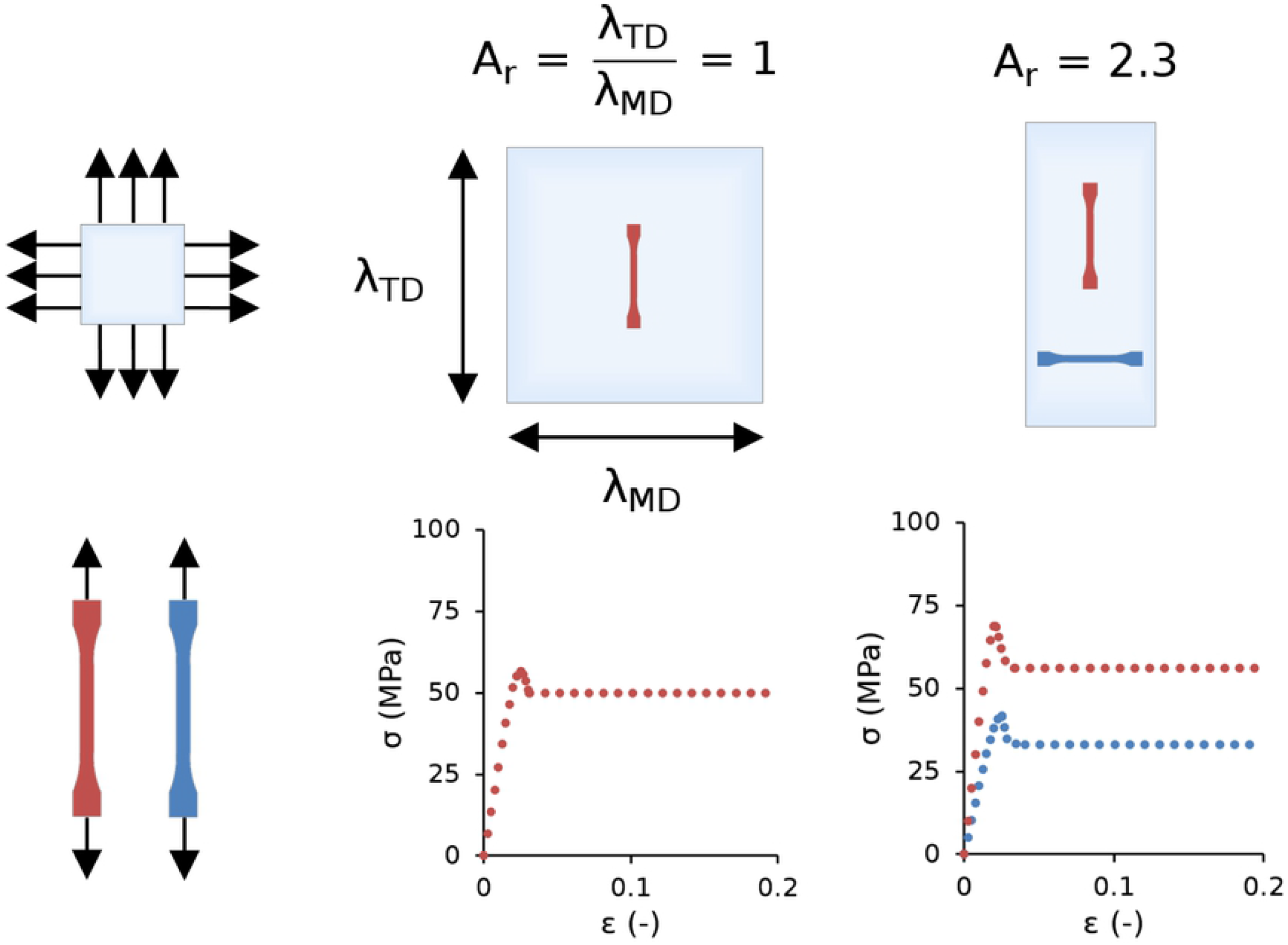
Schematic diagram showing experimental characterisation (from uniaxial tensile testing at 37 °C and 5 mm/min) for various aspect ratios (*A_r_*) of biaxially stretched *PLLA*.

In a follow-on study, Blair et al.^[43]^ varied *A_r_* and performed uniaxial tensile testing at various temperatures (20, 37 and 55 °C) and extension rates (1, 5 and 10 mm/min) — comparable conditions to those experienced by a stent.^[44]^ By tailoring *A_r_*, biaxially stretched sheets were processed with direction dependent (anisotropic) mechanical properties (Fig. 1). Results also showed that these mechanical properties were strongly dependent on temperature during uniaxial deformation, and not heavily dependent on extension rate. Empirical relations were developed that related *E* and *σ_Y_* to *A_r_* (Eq. 1–4) for 0.4 ≤ *A_r_* ≤ 2.3 and for a temperature of 37 °C (Fig. 2), and a transversely isotropic, rate-independent, elastic-plastic constitutive model was calibrated against uniaxial tensile test data. A simplified version of this model is proposed in the present study (Fig. 3) which neglects the softening following yield and assumes *PLLA* exhibits perfectly plastic behaviour, i.e. a change in strain causes no observable change in stress.

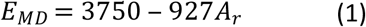

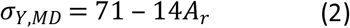

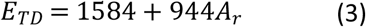

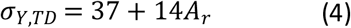

**Fig. 2.**
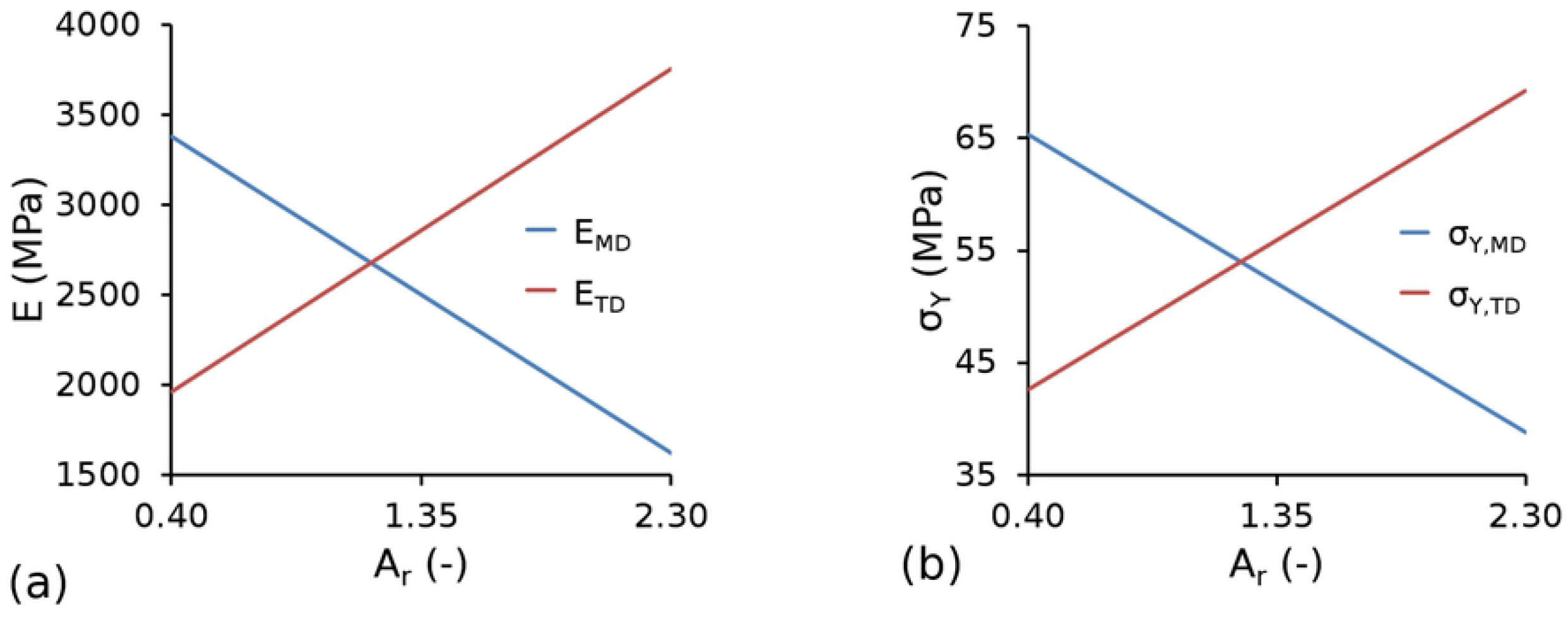
Graphical representation of constitutive equations showing (a) elastic modulus (*E*) and (b) yield strength (*σ_Y_*) in both the machine direction (*MD*) and transverse direction (TD) as a function of aspect ratio (*A_r_*).

**Fig. 3.**
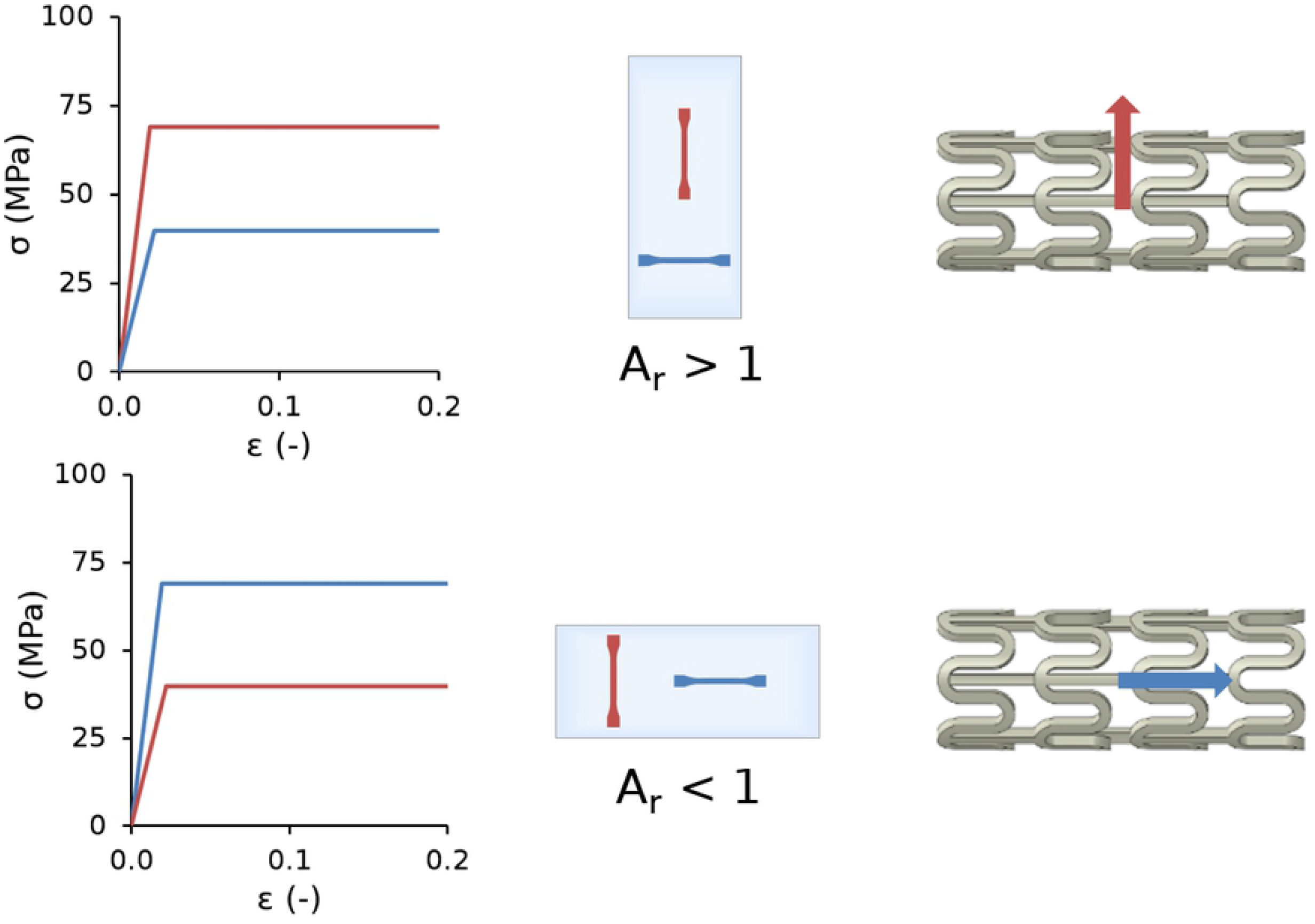
Schematic diagram showing the constitutive model stress-strain (*σ − ε*) curves for an *A_r_* > 1, which generates stents that are stiffer in the circumferential direction, and an *A_r_* < 1, which generates stents that are stiffer in the longitudinal direction.

Given that one of the most challenging aspects to overcome when designing polymer-based stents lies in the significantly lower radial stiffness compared to their metallic counterparts, it may be beneficial to process the stent such that it has a preferential circumferential orientation. An *A_r_* > 1 generated stent designs that are stiffer in the circumferential direction, whilst an *A_r_* < 1 generated stent designs that are stiffer in the longitudinal direction (Fig. 3).

### 2.2 Geometry parametrisation

The stent geometry used in the present study was based on a conventional open-cell stent design with straight bridges, using SolidWorks 2016 (Dassault Systèmes, France) to generate the three-dimensional model (Fig. 4). The stent was designed in the crimped state with two repeating unit cells used to represent the full-length stent geometry, thereby reducing computational cost. Parametric stent geometries were generated by varying the strut width (*w*), the strut thickness (*t*) and the strut length (*l*).

**Fig. 4.**
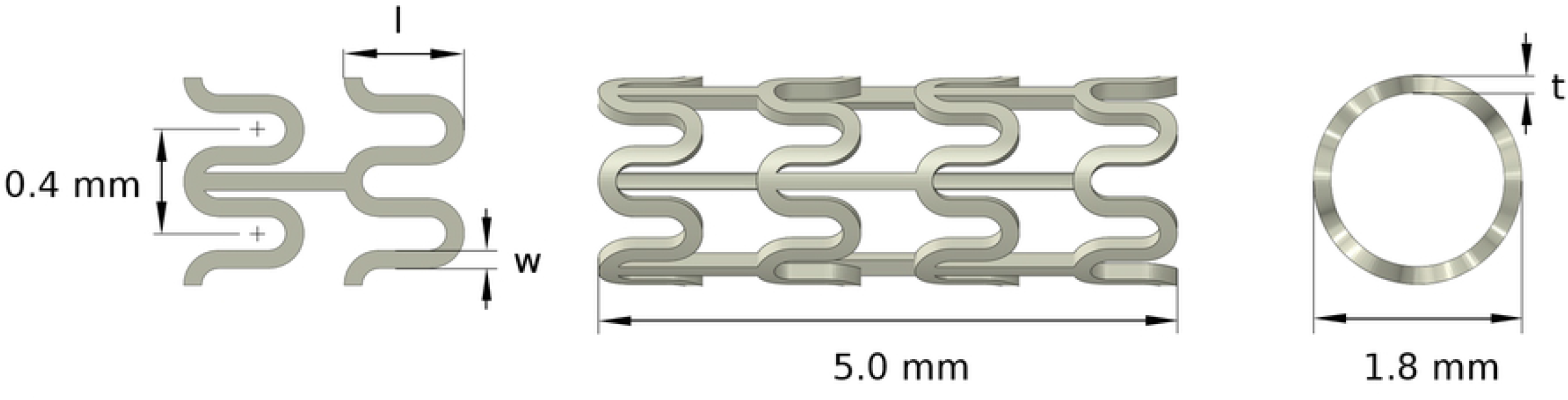
Geometry parameterisation in terms of strut width (*w*), strut thickness (*t*) and strut length (*l*).

### 2.3 Performance metrics

Four performance metrics were extracted for each stent design, based on the results of deployment and bench test simulations: (i) the cross-sectional area post-dilation (*CSA*), (ii) foreshortening (*FS*), (iii) stent-to-artery ratio (*SAR*) and (iv) radial collapse pressure (*RCP*). Initially, an idealised quasistatic expansion procedure was simulated in Abaqus/Standard 2016 (Dassault Systèmes, USA) using a displacement driven cylinder (meshed with S4R shell elements) and a deformable solid stent (meshed with C3D8R brick elements). The stent was designed in a pre-crimped state (Fig. 5a) and constrained in both the axial and tangential directions (with respect to a user-defined cylindrical coordinate system) via three nodes forming an equilateral triangle in the central section. A radial displacement was prescribed to all nodes on the cylinder increasing the stent diameter from 1.8 mm to 3.5 mm using the smooth-step amplitude definition within Abaqus, with tangential and axial displacement prohibited (Fig. 5b). Frictionless surface-to-surface contact was assumed, and self-contact was enabled for the stent. Following expansion, the cylinder was contracted during which the stent recoiled (Fig. 5c). The time-frame typically required for polymeric stent expansion approaches 1 min according to published guidelines from Abbott.^[45]^ However, given that a rate-independent material model is used, the time frame for expansion was reduced to 1 s.

**Fig. 5.**
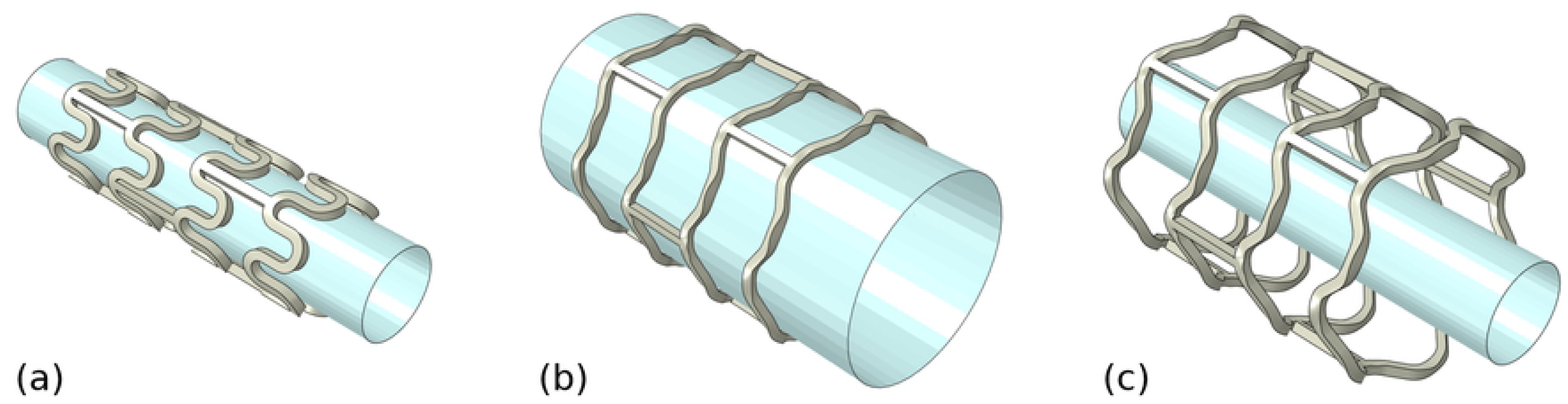
Finite element deployment simulation showing the stent in its (a) initial crimped state; (b) deployed (expanded) state and (c) final (recoiled) state.

The *CSA* following unloading was calculated based on the internal diameter of the stent (*D_unload_*) (Eq. 5) (Fig. 6a). During expansion, the opening of the strut hoops naturally cause the stent to contract in the axial direction (Fig. 6b). The *FS* of a stent was defined as the percentage reduction between the stent length in its crimped state (*L_initial_*) and the stent length following unloading (*L_unload_*) (Eq. 6). The *SAR* of the stent (Fig. 6c) was calculated as the ratio between the external surface area of the stent in its crimped state 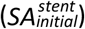 and the internal surface area of a compatible cylindrical artery (*SA^artery^*) (Eq. 7). The *RCP* of an expanded stent was evaluated through an additional virtual bench test in which eight rigid plates (meshed with R3D4 elements) (Fig. 6d) were radially contracted using a displacement driven process to produce 10% diameter loss. The *RCP* was calculated as the quotient of the average reaction force acting on the plates (*RF_ave_*) and the surface area of the stent post-recoil 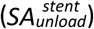 (Eq. 8). The smooth-step amplitude definition was used with frictionless surface-to-surface contact between the plates and the stent, and self-contact was enabled for the stent.

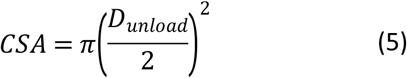

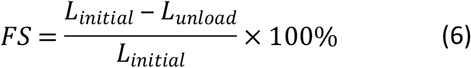

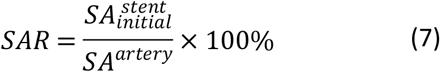

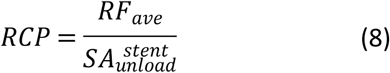

**Fig. 6.**
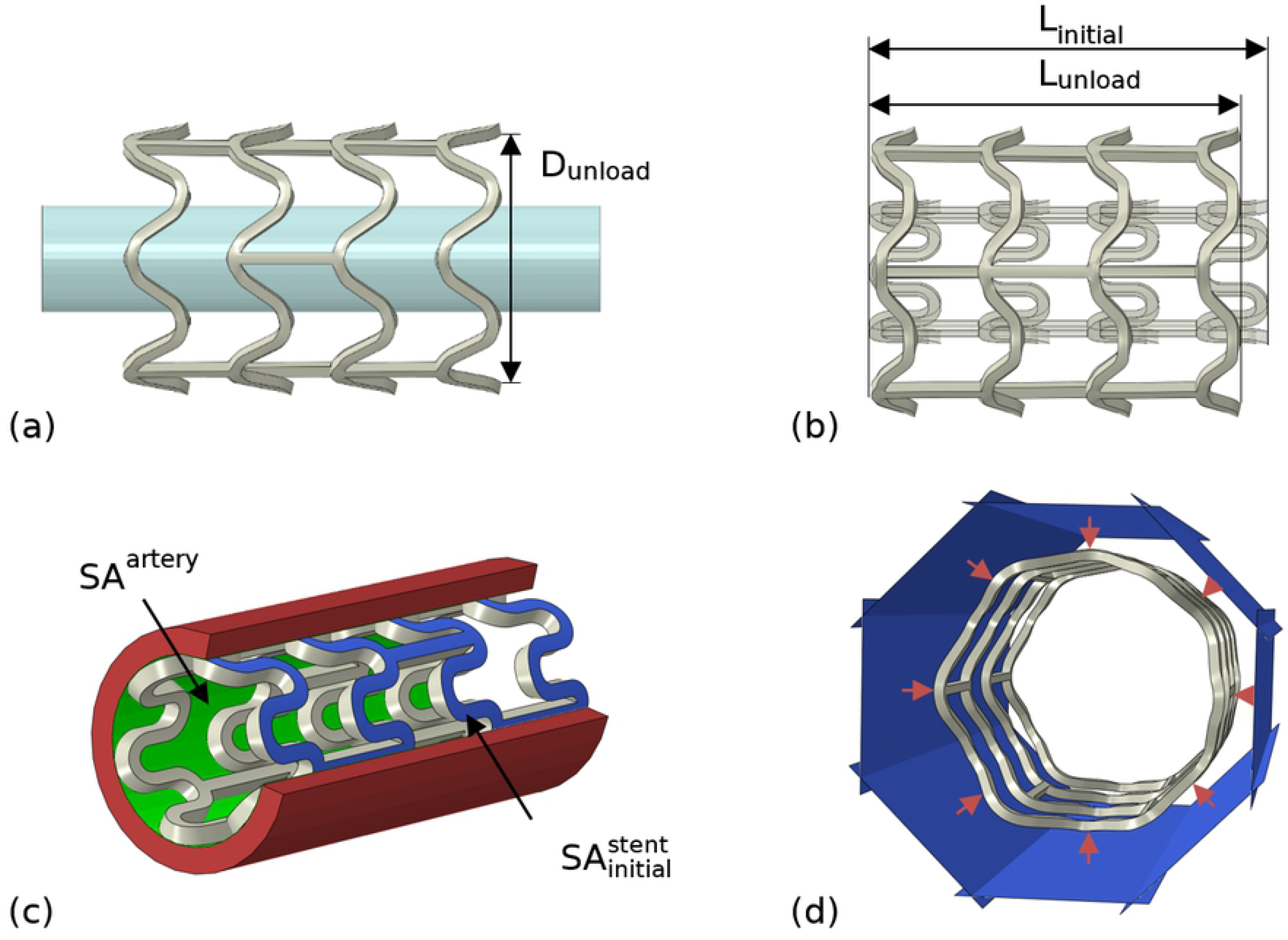
Schematic representations of tests for: (a) cross-sectional area (post-dilation), *CSA*; (b) foreshortening, *FS*; (c) stent-to-artery ratio, *SAR* and (d) radial collapse pressure, *RCP*.

### 2.4 Optimisation

The time required to perform the finite element simulations and calculate the performance metrics for a given parametric stent design exceeded 1 h using five parallel processors. At these time scales, global optimisation processes become computationally inefficient and the majority of optimisation studies tend to adopt surrogate modelling approaches.^[35]^ Hence, response surface methodology (*RSM*) was employed to provide an empirical correlation between processing and geometry parameters and the mechanical performance of the stent.

A design space was established using the limits for each of the design parameters (Table 2). The lower limit of *A_r_* generates stents that are stiffer in the axial direction whilst the upper limit generates stents that are stiffer in the circumferential direction. A lower limit of 100 μm was set for *w* and *t* to generate geometries that resembled a metallic stent, whilst an upper limit of 200 μm was set to generate geometries that resembled a polymeric stent. An upper limit of 1200 μm was set for *l* to avoid self-contact between neighbouring circumferential rings, whilst a lower limit of 900 μm was set to prevent excessive plastic deformation. A baseline design was generated by setting *A_r_, w, t* and *l* at the midpoint of their range.

**Table 2.** High and low levels for design parameters (*A_r_, w, t* and *l*).

| $A_r$ (-) | $w$ ($\mu\text{m}$) | $t$ ($\mu\text{m}$) | $l$ ($\mu\text{m}$) |
| --- | --- | --- | --- |
| 0.4 | 100 | 100 | 900 |
| 2.3 | 200 | 200 | 1,200 |

Initially, 40 design points that uniformly filled the design space were selected using an optimised Latin hypercube (LHC) sampling technique.^[46]^ Parametric stent designs and finite element models were automatically generated using a combination of Python (version 2.7.13; Python Software Foundation) scripting, SolidWorks 2016 (SolidWorks Corporation, USA) and the Abaqus CAE pre-processor. Deployment and bench testing simulations were performed in order to compute discrete values for each performance metric (*CSA, FS, SAR* and *RCP*). Multiple linear regression analysis was performed on the results using R (version 3.4.0)^[47]^ to provide an empirical correlation between each performance metric and design parameters. The Matplotlib (version 2.2.2) package^[48]^ was used to generate three-dimensional response surface plots to provide a qualitative, visual assessment of the results.

Following the RSM, multi-objective sequential least squares optimisation was performed in Python using the NumPy (version 1.14.2)^[49]^ and SciPy packages (version 1.2.0)^[50]^ to identify suitable options from non-dominated Pareto designs, i.e. a design that cannot be improved without degrading at least one of the other performance metrics. Each performance metric was normalised (scaled) to the same range [0,1], based on its minimum and maximum attainable values, attained through single objective sequential least squares minimisation. A single objective function (OF) was constructed (Eq. 9) that combines these normalised *CSA, FS, SAR* and *RCP* terms, with each parameter assigned an equal weighting. The intention of this optimisation was to minimise *FS* and *SAR* whilst maximising *CSA* and *RCP*. Hence, negative sign convention was adopted for *CSA* and *RCP* so that lower values for absolute and normalised performance metrics indicate better designs. An inequality constraint was imposed that prevented *RCP* dropping below 40 kPa (Eq. 10), which is commonly considered the minimum allowable collapse pressure for coronary stents.^[18]^ An additional inequality constraint was imposed that prevented *t* from exceeding the baseline value of 150 μm (Eq. 11).

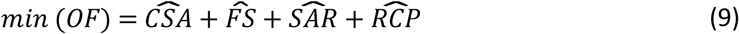

s.t.

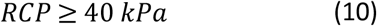

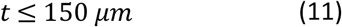

## 3. Results

### 3.1 Baseline geometry

The baseline stent design parameters and the respective performance metrics are shown in Table 3. Cross-sectional area (post-dilation) is difficult to measure *in vivo* and hence, there is limited published data. However, the baseline design recoiled by approximately 9% following dilation, which is comparable to commercial *PLLA BRS*.^[24]^ Given that the value of *t* is similar between the baseline design and a commercial stent, by extension, the *CSA* will also be comparable. The baseline stent design values for *SAR* and *FS* of 5.7% and 35.5%, respectively, are comparable to the upper end of the commercial *PLLA BRS* range.^[12,24]^ However, the baseline stent value for *RCP* of 20.9 kPa is approximately half of the minimum allowable collapse pressure for a coronary stent,^[18]^ thereby justifying the requirement for the present optimisation study.

**Table 3.** Baseline stent design parameters (*A_r_, w, t*, and *l*) and its respective performance metrics (*CSA, FS, SAR*, and *RCP*).

| $A_r$ (-) | $w$ ($\mu\text{m}$) | $t$ ($\mu\text{m}$) | $l$ ($\mu\text{m}$) | *CSA* ($\text{mm}^2$) | *FS* (%) | *SAR* (%) | *RCP* (kPa) |
| --- | --- | --- | --- | --- | --- | --- | --- |
| 1.35 | 150 | 150 | 1050 | -8.0 | 5.7 | 35.3 | -20.9 |

### 3.2 Response surface methodology

The four performance metrics (*CSA, FS, SAR* and *RCP*) were computed for each of the 40 design points (Table 4).

**Table 4.** Design parameters (*A_r_, w, t* and *l*) and respective performance metrics (*CSA, FS, SAR* and *RCP*) for each point considered under the optimised Latin hypercube sampling plan.

| *Design* | *A<sub>r</sub> (-)* | *w (μm)* | *t (μm)* | *l (μm)* | *CSA (mm<sup>2</sup>)* | *FS (%)* | *SAR (%)* | *RCP (kPa)* |
| --- | --- | --- | --- | --- | --- | --- | --- | --- |
| 1 | 0.90 | 101 | 191 | 1166 | -6.2 | 3.4 | 26.7 | -6.8 |
| 2 | 0.47 | 134 | 161 | 1001 | -8.1 | 6.8 | 30.9 | -18.9 |
| 3 | 0.52 | 119 | 154 | 1144 | -7.0 | 4.2 | 30.5 | -8.5 |
| 4 | 0.71 | 124 | 134 | 1039 | -7.6 | 5.5 | 29.9 | -13.1 |
| 5 | 2.23 | 146 | 169 | 1009 | -8.4 | 5.8 | 33.6 | -22.8 |
| 6 | 0.61 | 164 | 184 | 1016 | -8.4 | 8.5 | 37.2 | -35.3 |
| 7 | 0.57 | 179 | 156 | 956 | -8.8 | 10.0 | 38.4 | -39.9 |
| 8 | 1.90 | 189 | 189 | 1136 | -8.4 | 6.4 | 45.5 | -32.6 |
| 9 | 1.37 | 176 | 104 | 971 | -9.2 | 7.4 | 38.3 | -23.7 |
| 10 | 1.42 | 199 | 126 | 1084 | -8.5 | 7.2 | 45.9 | -27.7 |
| 11 | 1.94 | 151 | 121 | 1196 | -8.1 | 3.6 | 39.1 | -10.8 |
| 12 | 1.33 | 116 | 166 | 949 | -8.1 | 6.6 | 26.3 | -20.2 |
| 13 | 0.95 | 139 | 106 | 979 | -8.6 | 6.6 | 31.4 | -16.0 |
| 14 | 2.13 | 186 | 179 | 1046 | -8.7 | 7.2 | 42.4 | -36.8 |
| 15 | 1.80 | 169 | 146 | 994 | -8.7 | 6.8 | 37.6 | -29.4 |
| 16 | 1.23 | 161 | 176 | 904 | -9.0 | 9.5 | 33.8 | -45.3 |
| 17 | 1.52 | 129 | 144 | 1189 | -6.9 | 3.1 | 33.7 | -8.8 |
| 18 | 1.61 | 191 | 174 | 941 | -9.1 | 9.5 | 40.1 | -53.9 |
| 19 | 2.04 | 136 | 136 | 964 | -8.4 | 5.8 | 30.6 | -18.1 |
| 20 | 2.18 | 156 | 141 | 1076 | -8.9 | 5.1 | 37.2 | -18.6 |
| 21 | 1.09 | 194 | 111 | 911 | -9.0 | 10.5 | 39.6 | -35.0 |
| 22 | 1.18 | 141 | 124 | 1091 | -8.2 | 4.7 | 34.4 | -13.8 |
| 23 | 2.28 | 154 | 164 | 1174 | -7.8 | 4.1 | 39.1 | -14.5 |
| 24 | 1.28 | 196 | 196 | 1024 | -8.8 | 9.1 | 43.5 | -51.0 |
| 25 | 1.04 | 184 | 139 | 1114 | -8.2 | 6.6 | 43.9 | -24.7 |
| 26 | 1.99 | 126 | 114 | 1054 | -7.7 | 3.9 | 30.4 | -10.5 |
| 27 | 1.47 | 104 | 159 | 1069 | -7.0 | 3.8 | 25.7 | -9.2 |
| 28 | 1.56 | 131 | 199 | 1061 | -7.8 | 4.9 | 31.6 | -20.4 |
| 29 | 2.09 | 109 | 151 | 986 | -7.8 | 4.9 | 25.4 | -12.4 |
| 30 | 0.76 | 181 | 109 | 1129 | -8.0 | 6.7 | 43.8 | -17.4 |
| 31 | 1.75 | 149 | 194 | 934 | -8.8 | 8.0 | 32.3 | -38.0 |
| 32 | 0.66 | 171 | 129 | 919 | -8.8 | 9.7 | 36.0 | -31.6 |
| 33 | 0.99 | 106 | 116 | 926 | -7.8 | 6.1 | 23.9 | -12.9 |
| 34 | 0.42 | 159 | 131 | 1031 | -8.9 | 7.5 | 36.6 | -18.6 |
| 35 | 1.66 | 114 | 119 | 1121 | -6.9 | 3.2 | 28.9 | -6.9 |
| 36 | 1.85 | 111 | 186 | 1151 | -6.8 | 3.2 | 28.9 | -8.7 |
| 37 | 0.80 | 144 | 149 | 1181 | -7.2 | 4.6 | 37.0 | -12.1 |
| 38 | 1.71 | 166 | 101 | 1106 | -8.5 | 5.0 | 40.1 | -14.3 |
| 39 | 0.85 | 121 | 171 | 1099 | -7.2 | 4.7 | 30.2 | -12.7 |
| 40 | 1.14 | 174 | 181 | 1159 | -7.9 | 5.9 | 43.1 | -25.7 |

Multiple linear regression analysis (Table 5) was performed to generate constitutive equations that related each performance metric to the input parameters. A second-order model containing the intercept, main factors, two-factor interactions and quadratic terms (Eq. 12) was used for *CSA, FS, SAR* and *RCP*. Using the constants in Table 5, each model predicted, with approximately 99.7% confidence, that all values lie within the mean prediction plus or minus three standard deviations (Fig. 7). Model quality is assessed in Fig. 8, in which the performance metrics were predicted for a given set of design parameters using the statistical model (Eq. 12), and compared to their corresponding actual (measured) values extracted from finite element simulations. Linear behaviour was observed for *CSA, FS, SAR* and *RCP*, with the statistical models achieving R-squared (R^2^) values of 0.950, 0.996, 0.999 and 0.996, respectively.

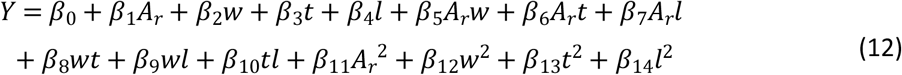

where Y denotes the predicted response for a given performance metric, i.e. *CSA, FS, SAR* and *RCP*.

**Table 5.**
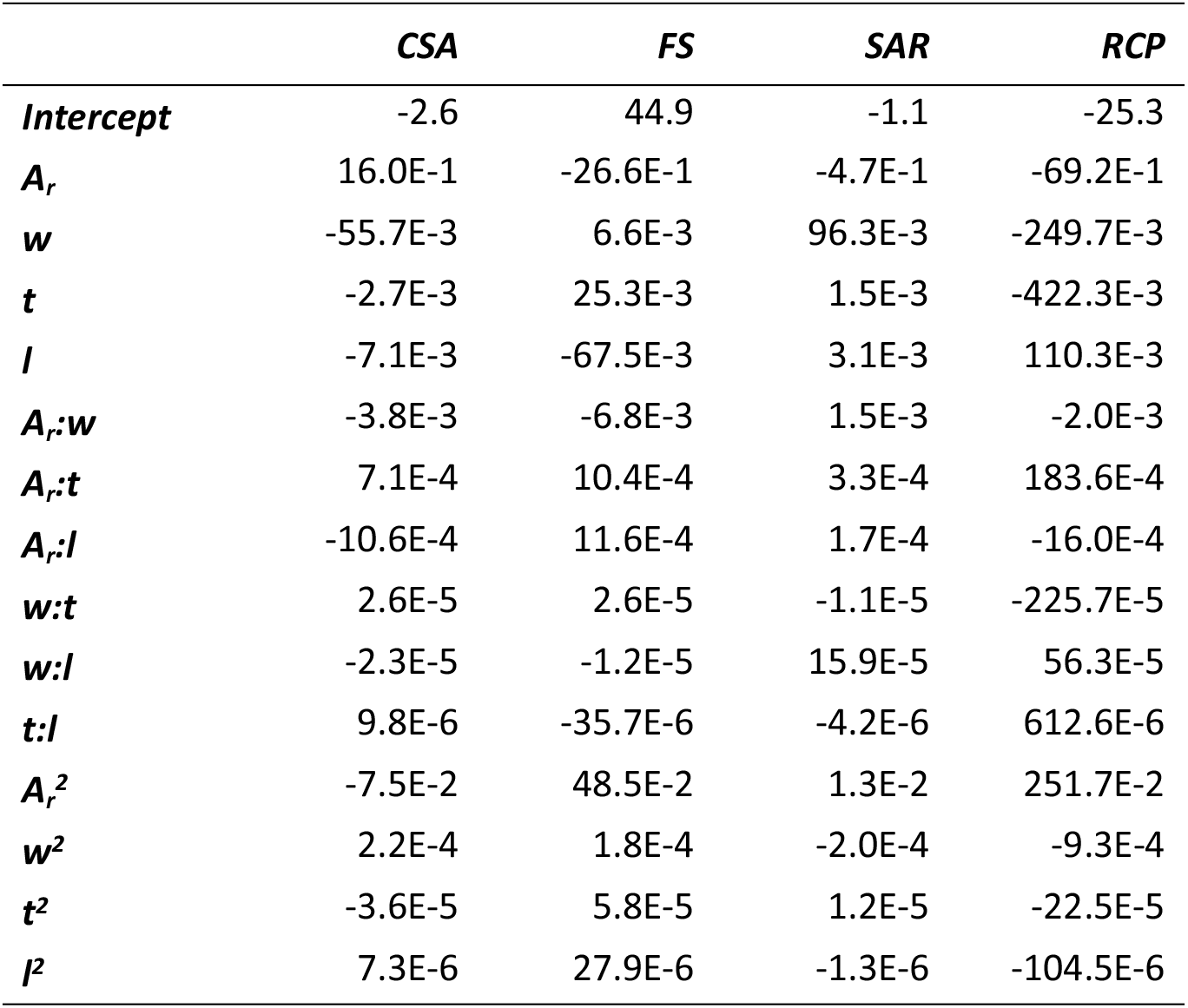
Statistical model coefficients for *CSA, FS, SAR* and *RCP*.

**Fig. 7.**
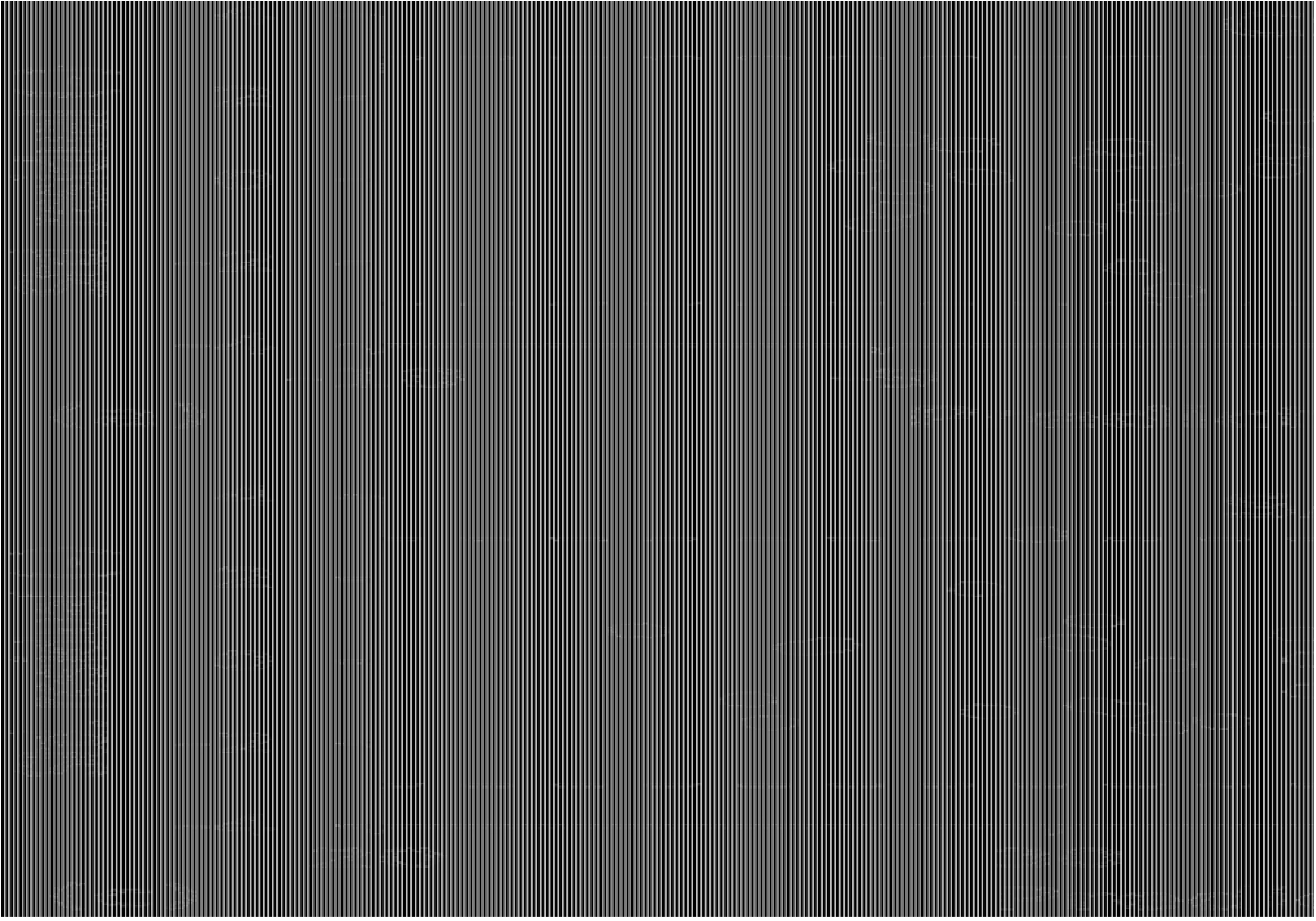
Standardised residual vs. predicted response using the statistical model in Eq. *Error! Reference source not found*. for (a) *CSA*; (b) *FS*; (c) *SAR* and (d) *RCP*.

**Fig. 8.**
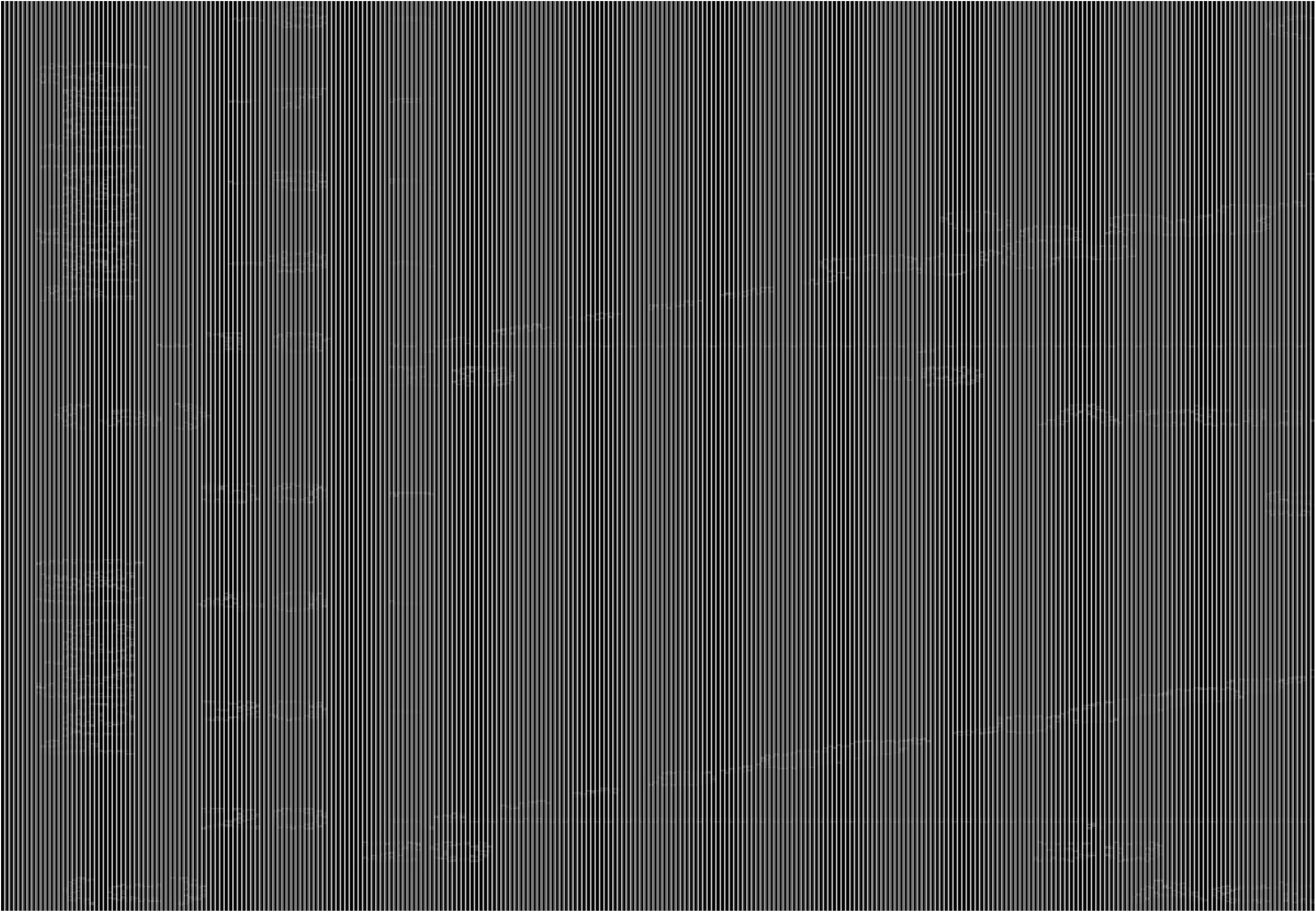
Predicted response using the statistical model in Eq. *Error! Reference source not found*. vs. actual (measured) response from finite element simulations for (a) *CSA*; (b) *FS*; (c) *SAR* and (d) *RCP*.

A comparison of absolute t-values (for coefficients) from multiple regression analyses for each performance metric is shown in Fig. 9a–d. Main factors, two-factor interactions and quadratic terms are considered statistically significant (*p* < 0.05) if their absolute t-value lies above the dashed line.

**Fig. 9.**
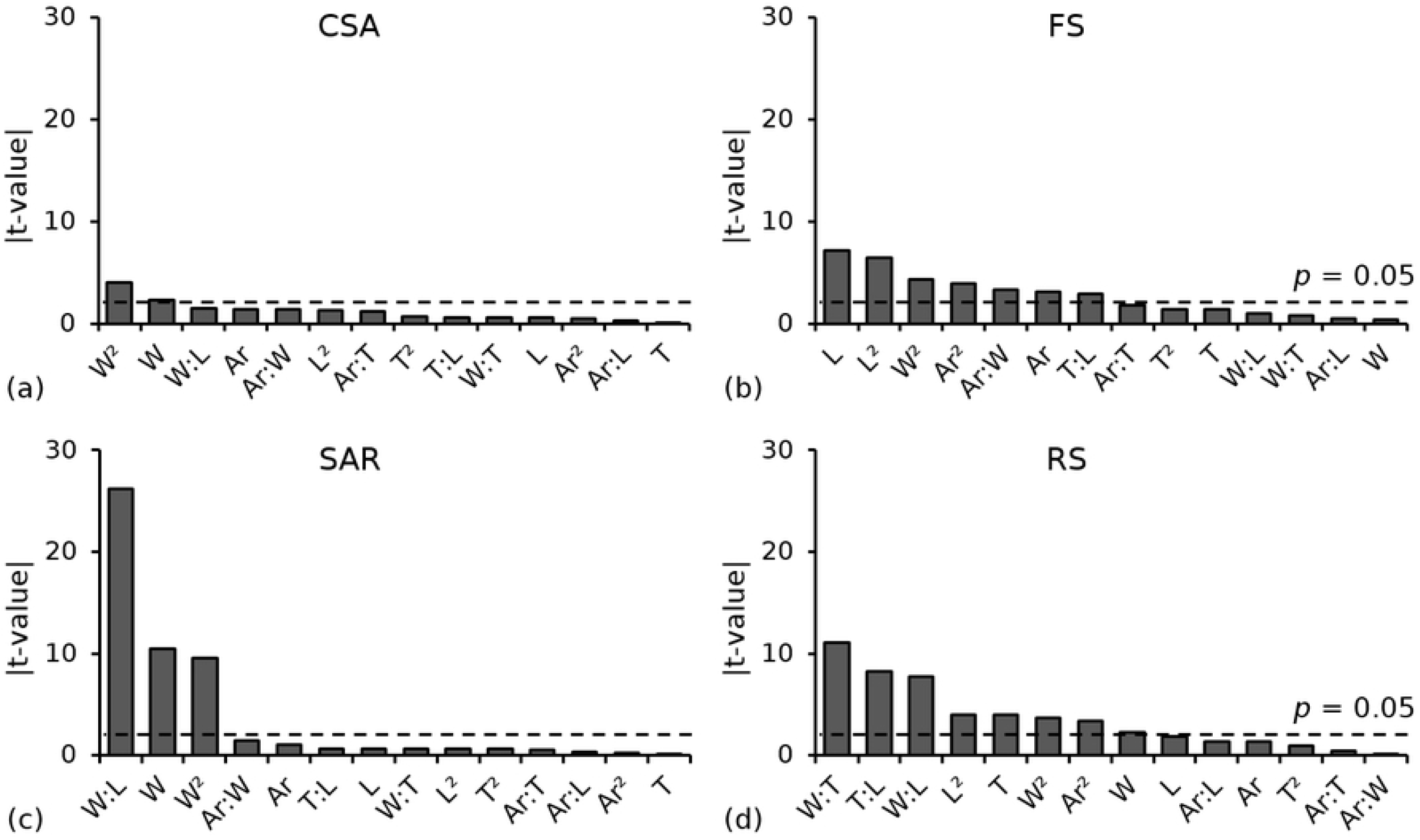
Comparison of absolute t-values (for coefficients) from multiple regression analyses highlighting significant (p < 0.05) main factors and two-way interactions for (a) *CSA*; (b) *FS*; (c) *SAR* and (d) *RCP*.

Response surfaces were plotted for all two-way interactions (Fig. 10), which highlight the combined influence of any two design parameters (*A_r_, w, t* or *l*) on each performance metric (*CSA, FS, SAR* and *RCP*). For each response surface, the performance metric was plotted against two dependent design parameters whilst the remaining two independent parameters were held constant at their baseline (midpoint) value. For each response surface, moving from the purple region to the yellow region indicates an improvement.

**Fig. 10.**
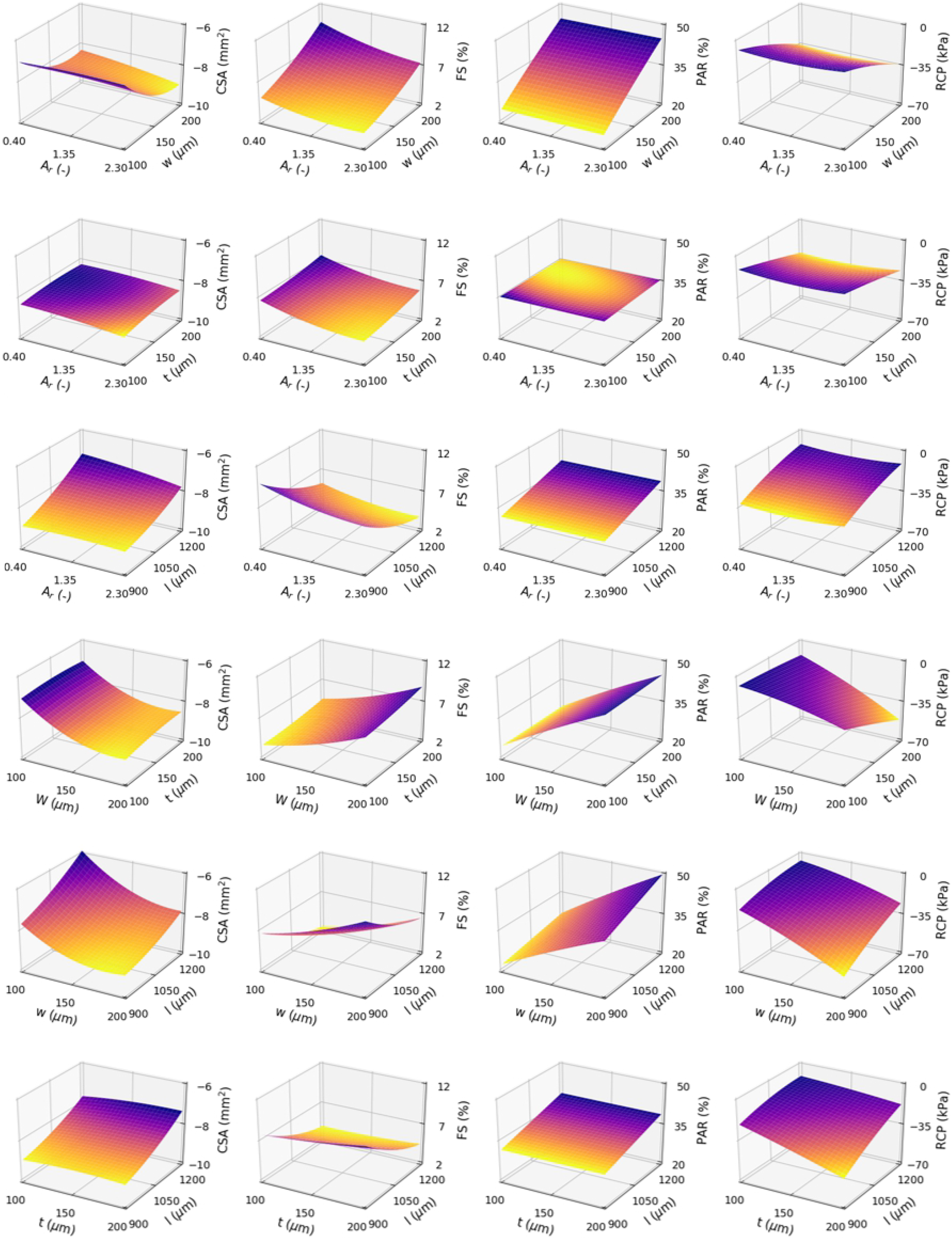
Response surfaces highlighting the combined influence of any two design parameters (*A_r_, w, t* or *l*) on each performance metric (*CSA, FS, SAR* and *RCP*). For each response surface, the remaining two (independent) design parameters are held constant at their baseline value.

The Pareto fronts (Fig. 11) highlight the trade-offs between each set of performance metrics, with better designs lying towards the bottom left corner. Trade-offs were observed for *CSA vs. FS, CSA vs. SAR, FS vs. RCP* and *SAR vs. RCP*, whilst no trade-offs were observed for *CSA vs. RCP* or *FS vs. SAR*. Trade-offs occurred as a result of conflicting requirements for stent design, i.e. geometric and/or material parameters that improve one metric often negatively affect at least one of the other metrics.

**Fig. 11.**
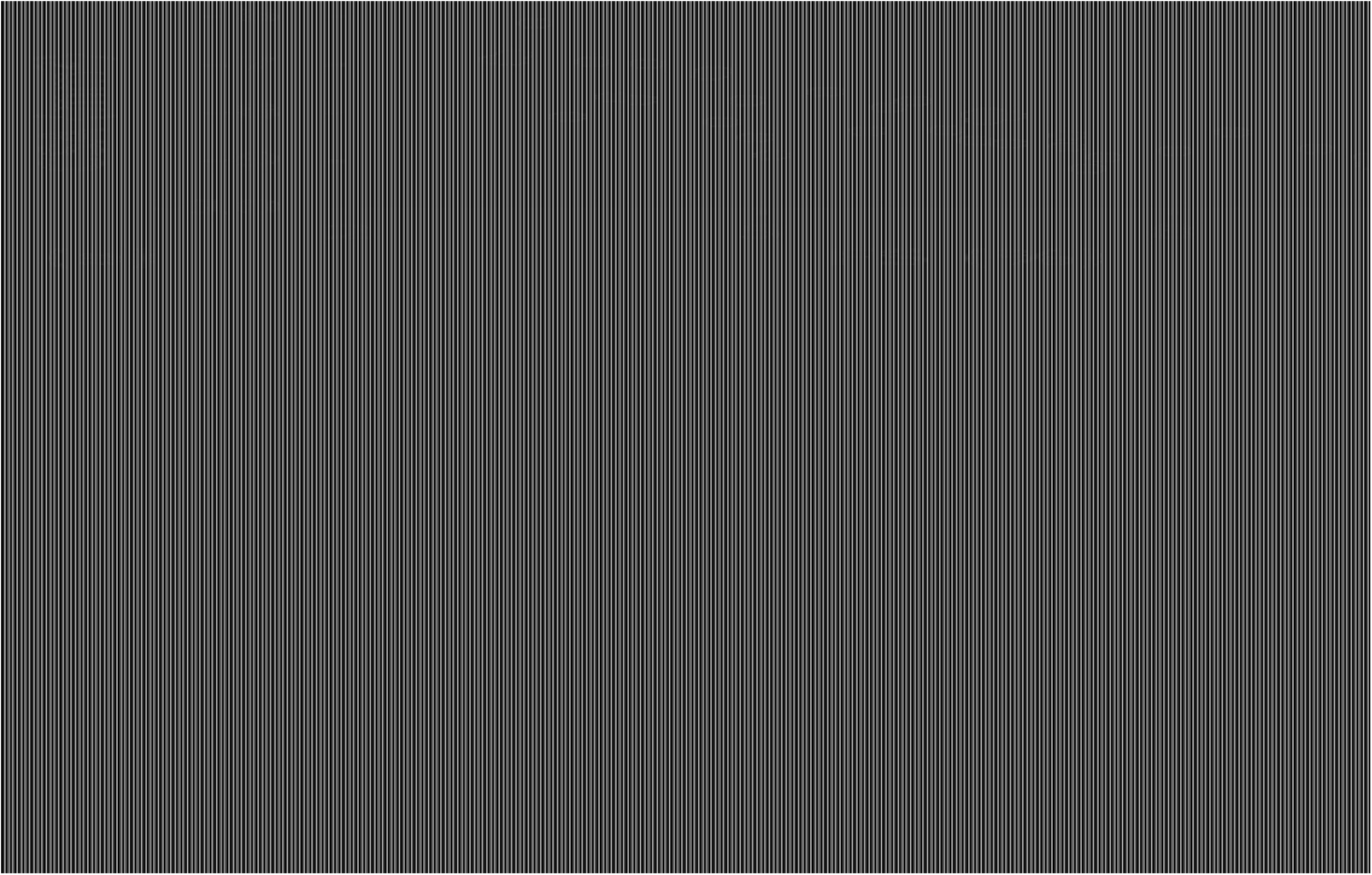
Trade-off curves for all permutations of the four performance metrics: (a) *CSA* vs. *FS*; (b) *CSA vs. SAR*, (c) *CSA vs. RCP* and (d) *FS vs. SAR*, (e) *FS vs. RCP* and (f) *SAR vs. RCP*.

### 3.3 Optimisation

To construct a single dimensionless objective function, each performance metric was normalised (scaled) to the same range [0,1] based on its minimum and maximum attainable values (Table 6), attained using least squares minimisation (Eq. 13).

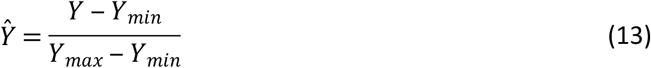

where *Ŷ* and *Y* denote the predicted normalised and absolute responses, respectively, for a given performance metric, whilst *Y_min_* and *Y_max_* denote the minimum and maximum attainable values.

**Table 6.**
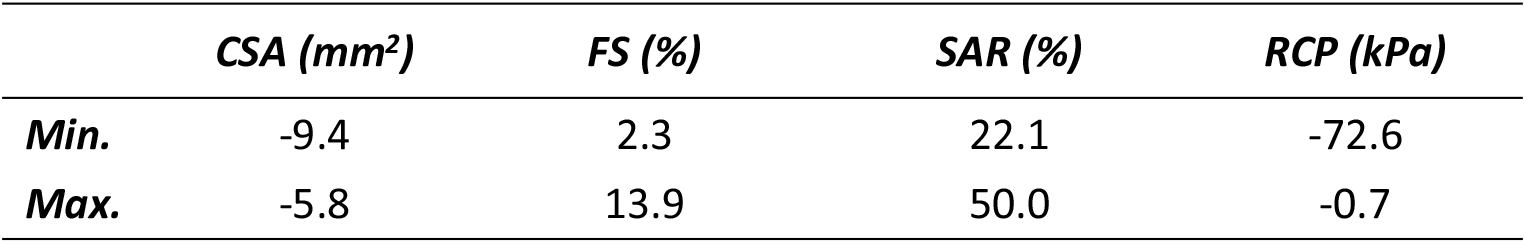
Minimum and maximum values for each performance metric (*CSA, FS, SAR* and *RCP*).

Multi-objective optimisation produced a stent design superior to the baseline with *t* = 150 μm and *w* = 173 μm (Table 7), which are lower than some commercial polymeric stents,^[12]^ whilst meeting the minimum allowable collapse pressure.^[18]^ A comparison between the baseline design and the optimised design is shown in Fig. 12, in which each performance metric has been normalised. The *RCP* of the optimal design is approximately twice that of the baseline design with a less than 1% increase in *SAR*. The *CSA* increased by 14% and whilst *FS* increased, a value of 8% is comparable to stents in commercial use.^[51]^

**Fig. 12.**
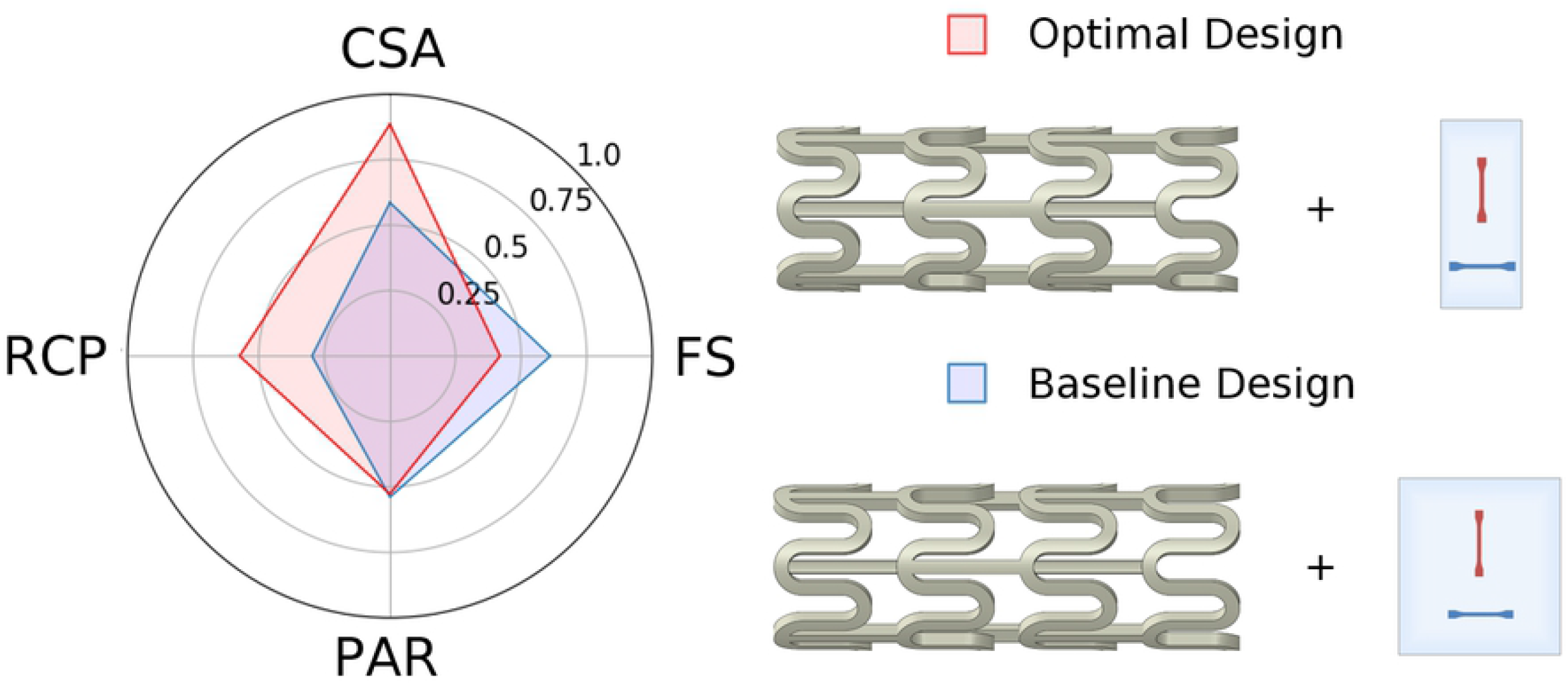
Visual comparison of normalised performance metrics and design parameters between the baseline design and the optimal design.

**Table 7.** Comparison between baseline (base.) and optimal (opt.) stent designs highlighting design parameters and their respective performance metrics.

| | $A_r$ (-) | $w$ ($\mu m$) | $t$ ($\mu m$) | $l$ ($\mu m$) | $CSA$ ($mm^2$) | $FS$ (%) | $SAR$ (%) | $RCP$ (kPa) |
| --- | --- | --- | --- | --- | --- | --- | --- | --- |
| **Base.** | 1.35 | 150 | 150 | 1050 | -8 | 5.7 | 35.3 | -20.9 |
| **Opt.** | 2.3 | 173 | 150 | 900 | -9.1 | 8 | 35.7 | -40 |

## 4. Discussion

This study proposes a multi-objective optimisation framework that considers the combined effect of the biaxial stretching processing history and the geometric configuration when optimising the short-term (pre-degradation) mechanical performance of a *PLLA* coronary stent. Given that the ideal stent must fulfil a range of conflicting technical requirements, a multi-objective optimisation process that offers compromises between key performance metrics was conducted to develop a polymeric stent that offered improved performance relative to a baseline design for the same strut thickness (150 μm). Performance trade-offs were observed (Fig. 11) and may be explained using the absolute t-value comparisons for coefficients (Fig. 9a–d) and the response surface interaction plots for each performance metric (Fig. 10). The absolute t-value comparisons for coefficients highlight statistically significant (*p* < 0.05) factors for each performance metric whilst the response surface interaction plots provide a visual aid in understanding the interdependent effect between two factors on a given performance metric.

### 4.1 Cross-sectional area *vs*. foreshortening

The trade-off between *CSA* and *FS* was primarily due to the conflicting requirements for w and *l*. Cross-sectional area was most strongly affected by w and *w*^2^ (Fig. 9a), whilst *FS* was most strongly affected by *l* and *l*^2^ (Fig. 9b). Increasing *w* improved *CSA* as a wider strut increased plastic deformation in the hoops and reduced radial recoil, which is in agreement with the findings of Pant et al.^[40]^ Furthermore, the presence of a significant (*p* < 0.05) quadratic effect (*w*^2^) in the model suggested a curvilinear relationship between *CSA* and *w*. This was evident from the interaction plots in which *w* was plotted as one of the dependent variables (Fig. 10). A convex relationship was observed between *CSA* and *w*, i.e. *CSA* improved as *w* increased but with diminishing returns. Decreasing *l* further improved *CSA* and was evident from the interaction plot between *w* and *l*. By increasing *w* from 100 μm to 200 μm and decreasing *l* from 1,200 μm to 900 μm, *CSA* improved by approximately 53%. However, this change caused an undesirable increase in *FS* from 3% to 11%. In contrast to the requirements for *CSA*, narrow, long struts were ideal for reducing *FS*, as the struts deformed less to achieve an equivalent level of plastic strain, thereby reducing the level of axial contraction. This is in agreement with Li et al.^[39]^ who acknowledged the contrasting requirements for l, based on the observed trade-off between recoil and *FS*. Strut thickness has the weakest effect on *CSA* — whilst a higher value of t reduced the degree of radial recoil post-inflation, it was not offset by the reduced *CSA* (as a result of the thicker struts) pre-inflation. In general, it was beneficial to design the stent such that it is stiffer in the circumferential direction (higher *A_r_*) as *FS* improved without negatively affecting *CSA*. Hence, a lower value of *l* and *A_r_* were desirable.

### 4.2 Cross-sectional area vs. stent-to-artery ratio

The trade-off between *CSA* and *SAR* was primarily due to the conflicting requirements for *w*. Although high values of *w* improved *CSA*, a wider strut increased the surface area of the stent which negatively affects *SAR*. Low values of *l* were correlated with improved *CSA*, and were also correlated with improved *SAR* as, intuitively, a shorter strut reduced the surface area of the stent. The interaction between *w* and *l* had the strongest effect on *SAR* (Fig. 9c) and was evident from the response surface plot (Fig. 10). Stent-to-artery ratio was unaffected by *t* and *A_r_* and hence, it was beneficial to design the stent with high values of *A_r_* and *t* as these parameters improved *CSA*. High values of *A_r_* and *t*, combined with a low value of *l* are ideal for improving both *CSA* and *SAR*. By holding each of these design parameters constant at their optimal limits and increasing *w* from 100 μm to 200 μm, *CSA* improved by approximately 20%. However, *SAR* had an undesirable increase from 22% to 40%, which is significantly higher than the *SAR* for both polymer and metallic stents in clinical practice, and may contribute to increased levels of thrombosis.^[12,13]^

### 4.3 Foreshortening *vs*. radial collapse pressure

The trade-off between *FS* and *RCP* was primarily due to the conflicting requirements for *w, t* and *l*. Radial collapse pressure was most strongly affected by the interactions between *w* and t, *w* and *l* and *t* and *l*, with each interaction considered statistically significant (*p* < 0.05) (Fig. 9d). The response surface plots for each of these interactions (Fig. 10) showed that *RCP* improves with high values of *t* and w, combined with low values of *l*. This combination of parameters tended to induce higher levels of plastic deformation in the strut hoops. By increasing *w* and *t* from 100 μm to 200 μm and decreasing *l* from 1,200 μm to 900 μm, *RCP* improved from 8.8 kPa to 70 kPa, meeting the minimum allowable collapse pressure of 40 kPa.^[18]^ However, this change caused an undesirable increase in *FS* from 2.5% to 12%. In general, *A_r_* did not strongly affect *RCP* and was not considered statistically significant (*p* > 0.05). However, given that a higher *A_r_* improved *FS*, it was beneficial to design the stent such that it is stiffer in the circumferential direction.

### 4.4 Stent-to-artery ratio vs. radial collapse pressure

The trade-off between *SAR* and *RCP* is similar to the trade-off observed between *SAR* and *CSA*, and is primarily due to the conflicting requirements for *w*. High values of *A_r_* and *t*, combined with a low value of *l* are ideal for improving both *RCP* and *SAR*. By holding each of these design parameters constant at their optimal limits and increasing *w* from 100 μm to 200 μm, *RCP* had a more than three-fold increase. However, SAR had an undesirable increase of approximately 80%.

### 4.5 Limitations

In this study, stent geometries were based on a conventional open-cell design with straight bridges, which has proved ideal for metallic drug-eluting stents. However, this does not guarantee compatibility when using a polymer such as *PLLA* as the platform material, given that it exhibits an entirely different stress-strain response. Modifying the bridge geometry, strut cross-section and hinge profile have all been shown to influence the mechanical performance of stents^[40,52]^ and the inclusion of these parameters may permit the evaluation of unconventional (or unorthodox) geometries that are better suited to polymeric stents. In addition to increasing the number of design parameters, the inclusion of a stenosed artery into the finite element model would permit additional performance metrics to be evaluated. Modelling the expansion of a stent in a stenosed artery could provide an indication of high risk areas in the stented region and may also be used to evaluate the stent’s susceptibility to fracture. However, increasing the number of design parameters and performance metrics will increase the computational cost and complexity of the optimisation. Given that the performance metrics and design parameters evaluated within the present study were considered most critical based on the literature reviewed, any alternatives should be evaluated as additions rather than replacements. Finally, there is limited information in literature on clinically acceptable values for performance metrics such as foreshortening and stent-to-artery ratio. Identification of operational limits for these metrics is essential, as these limits can be used as constraints for the multi-objective optimisation procedure to tailor stent designs for a particular lesion or patient geometry, suggesting an area for future research.

## 5. Conclusion

An optimisation framework has been proposed that considers the combined effect of the biaxial stretching processing history and the geometric configuration when optimising the mechanical performance of a *PLLA* coronary stent. Response surface methodology combined with multi-objective optimisation produced an optimal *PLLA* stent design that offered improved performance relative to a baseline design for the same strut thickness (150 μm). The effects of each of the design parameters (*A_r_, w, t* and *l*) on individual performance metrics (*CSA, FS, SAR* and *RCP*) have been quantified and compared. For each of the design parameters, a main factor or two-way interactions term had a statistically significant (*p* < 0.05) effect on at least one of the performance metrics. Pareto fronts highlighted that a change in one design parameter that improves one metric often leads to a compromise in at least one of the other metrics with trade-offs observed for *CSA vs. FS, CSA vs. SAR, FS vs. RCP* and *SAR vs. RCP*. In summary, this study addresses key limitations in polymeric stent design and the methodology that could be applied in the development of high stiffness, thin strut polymeric stents that contend with the performance of their metallic counterparts.

## Acknowledgements

The authors wish to acknowledge funding from the Engineering and Physical Sciences Research Council (EPSRC) (S3804ASA) and the Marie Skłodowska-Curie Research and Innovation Staff Exchange (RISE), grant agreement 691238. The authors also wish to acknowledge the research institutions involved in the Bi-Stretch-4-Biomed collaborative RISE project (California Institute of Technology, University of Warwick and ENEA: Italian National Agency for New Technologies, Energy and Sustainable Economic Development).

